# Effects of an experimental contamination by methyl-mercury and glyphosate on biological and otolith shape parameters of a tropical Siganidae

**DOI:** 10.64898/2026.11.25.769091

**Authors:** Fabien Morat, Nelly Wabete, Chloe Lebrun, Florian Geslin, Maxime Millot, Florence Antypas, Paco Bustamante, Valeriano Parravicini, Emmanuel Tessier, Yves Letourneur

**Author notes:** Corresponding author: Fabien Morat, PSL Université Paris: EPHE-UPVD-CNRS, UAR 3278 CRIOBE, Université de Perpignan, 52 Avenue Paul Alduy, 66860 Perpignan, France. **Funding Information**: TONIC project was funded by the French National Research Agency (ANR-21-CE32-0008).

## Abstract

This study examines the responses of a tropical herbivorous fish, the golden-lined spinefoot *Siganus lineatus*, to several experimental concentrations of two major contaminants, i.e., methyl-mercury (MeHg) and glyphosate. These contaminants were introduced *via* the fish feed, which itself contained varying concentrations. After eleven weeks of growth in the tanks, the fish were euthanised and two sets of parameters were investigated. The first type focuses on usual biological aspects (body condition, growth, energy storage, etc.), whilst the second type of parameters focuses on *lapillus* and *sagitta* otoliths (including various shape metrics). Overall, no significant differences were detected in the biological parameters during this experiment, except for the hepatosomatic index, which was lower in fish exposed to high concentrations of glyphosate. These results contradict the intuitive assumption that MeHg and glyphosate have effects on most biological parameters, as is often described in the literature. Conversely, the effects of contaminants are more noticeable in the otoliths, with significant differences in the shape of the *lapilli* and *sagittae* between fish that were non-contaminated or only slightly contaminated and those that were more heavily contaminated. The *sagittae* were more sensitive to MeHg, whilst the *lapilli* were more sensitive to glyphosate. As otoliths play a role in the sensory function of fish (orientation and positioning in the water column, perception of sound, etc.), our results suggest that high concentrations of MeHg and/or glyphosate in the natural environment could have harmful effects on the physiological and behavioural responses of this species.

## Introduction

The fish inner ear represents a combined stato-acoustic, multi-sensory organ involved in three different functions: (i) the semi-circular canals detect the angular accelerations resulting from head – body rotation, (ii) the otoliths detect linear accelerations resulting from swimming and body tilts, and (iii) they detect the sound (Popper and Combs, 1980; Popper *et al*., 2005; Schulz-Mirbach *et al*., 2019). Three otolith pairs are found in fish inner ear: the *lapilli* in utricle, the *sagittae* in saccule and the *asterici* in lagena. The role of these otoliths in these functions is still uncertain; the ‘mixed function hypothesis’ proposes that each otoliths are implicated in both vestibular and auditory senses (Popper and Fay, 1993; Schulz-Mirbach *et al*., 2019), whereas a second hypotheses proposes a ‘division of labour’ with the utricle implicated in vestibular function and the saccule and lagena in sound detection (Inoue *et al*., 2013; Schulz-Mirbach *et al*., 2019). For example, Yao *et al*. (2016) suggest that both utricle and saccule of zebra fish (*Danio rerio*) larvae are involved in audition (increase of microphonic threshold to high frequencies when saccule was removed and for low frequencies when utricle was removed). In the other hand, Roberts *et al*. (2017) showed that the normal swimming behaviour of mutant zebrafish larvae (dorsal-up postural) is observed only when the utricle are formed, suggesting its function on gravitational sense.

Otoliths are acellular calcium carbonate structures, which grow through the life time (Campana and Neilson, 1985). They are constituted by daily formation of alternating mineral-rich and protein-rich bands (Jolivet *et al*., 2008; Thomas and Swearer, 2019) and are metabolically inert (Campana, 1999; Campana and Thorrold, 2001). Associated with growth increments (daily and annual) that they formed, they provide information about individual age and life history (Campana, 2005). The otoliths were used in a wide range of studies such as species identification (Aguirre and Lombarte, 1999; Morat *et al*., 2012) and predators diet determination (Lilliendahl and Solmundsson, 2006; Byrd *et al*., 2020) as their shape is species-specific; age and growth assessment (e.g., Cardinale *et al*., 2004; Morat *et al*., 2020; Logez *et al*., 2021; Andrialovanirina *et al*., 2023), stock assessment (Friendland and Reddin, 1994), and to determine the fish migration or population connectivity (e.g., Vasconcelos *et al*., 2008; Morat *et al*., 2014; Bounket *et al*., 2019). Within a species, the otolith shape can vary according both environment and genetic factors (Vignon and Morat, 2010). The ecomorphological link hypothesis of otolith showed that the shape varies with age, size and ontogeny (Cardinale *et al*., 2004; Hüssy, 2008; Bounket *et al*., 2019), with the ecological gradient (driven by depth, temperature, etc.) and was used for discriminate local fish population according to the environmental characteristics (Campana and Casselman, 1993; Friendland and Reddin, 1994; Mérigot *et al*., 2007; Morat *et al*., 2012). For example, Bounket *et al*. (2019) showed that the *lapillus* shape of juveniles of chub *Squalius cephalus* varied with ontogeny and was stabilized above 120 days post-hatching. Vignon (2012) showed that the *sagitta* shape of the common bluestripe snapper *Lutjanus kasmira* varied with ontogeny in relation with fish growth and size, but this shape was changed with an important shift of coral reef habitat during the fish life cycle (estuarine to channel or outer slope).

Studies on the influence of otolith morphology on physiological functions are scarce (Schulz-Mirbach *et al*., 2019). The otolith crystallisation can change the sound perception; Oxman *et al*. (2007) showed that fish with abnormal otolith (i.e., vaterite instead of aragonite) had a decrease of sound sensitivity possibly related to lower vaterite otolith density and to a fewer sensory hair bundles in the saccular epithelium. Vignon and Aymes (2020) evidenced that juveniles of brown trout *Salmo trutta* with vaterite otoliths had an altered escape kinematic in response to sound stimuli. The size and shape of the *sulcus acusticus* (a groove along the surface of the *sagitta*) can also change the sound perception (Tuset *et al*., 2016; Schulz-Mirbach *et al*., 2019). Indeed, the *sulcus acusticus* of *sagitta* and the *gibbus maculae* (more or less protuberance in ventral face) of *lapillus* are in contact with saccule and utricle maculae, respectively (Assis, 2005; Schulz-Mirbach *et al*., 2019). The macula, the sensory epithelium, comprises the hair cells bundles implicated detection of otoliths motion and so the auditory mechanism (Pisam *et al*., 1998; Popper *et al*., 2005). Popper and Coombs (1982) showed that the *ostium* size (anterior part of the *sulcus acusticus*) was correlated with the proportion of the hair cells implicated in the detection of directional acoustic stimuli (Popper and Fay, 1993).

Coral reefs undergo natural and anthropogenic disturbances; Burke *et al*. (2011) estimated that 20-25% of coral reefs were impacted by contaminants. This assessment is however likely to be underestimated; for instance, Wejieme *et al*. (2025a, b) demonstrated that all the contaminants they studied were found in each coral reef fish individual at the whole SW Pacific scale. The environmental contamination can arise from various origins such as volcanic eruptions, industrial and wastewater discharges, mining activities, agricultural activities, or soil leaching that release into the environment contaminants such as pesticides and metallic trace elements. These contaminants, resistant to degradation, can be dispersed at long distance (Phillips, 1995) and can be toxic at low concentrations for organisms and humans.

Pesticides are known to impact growth, physiology, reproduction, immune response, etc. of aquatic organisms. In fish, they can alter the embryonic development, the larval growth (e.g., Shahjahan *et al*., 2017; Hasan *et al*., 2022); and they induce histopathological and haemato-biochemical alterations (for large review see Rohani, 2023). In the large group of pesticides, the glyphosate is a large spectrum herbicide active at low doses, considered as the most widely used herbicide in the world (Tarazona *et al*., 2017). Danger of glyphosate for mammals are still debated (Tarazona *et al*., 2017) but impacts has been observed for several aquatic organisms including fish. Thus, direct glyphosate exposure can increase larvae mortality and morphological alteration on the zebrafish *Danio rerio* and the silverside *Odontesthes humensis* (Zebral *et al*., 2017; Lanzarin *et al*., 2023) and juvenile of brown trout *Salmo trutta fario* (Drechsel *et al*., 2024). On the zebrafish, it can induce changes in hatching success and timing (delay, acceleration or inhibition), functional heart changes, neurotoxicity and behavioural changes, induce oxidative stress, gene expression changes, induce apoptosis, induce endocrine disruption and reproduction impairment, induce energy and metabolism disruption, and immunotoxicity (Lanzarin *et al*., 2023). Glyphosate exposure can also alter organs such as gills (proliferation of filament cell, fused lamellae, etc.), liver (necrosis) and kidney (hyaline droplets and pyknotic cells) on various species (Rohani, 2023). Other studies on characin *Leporinus obtusidens* and *Oreochromis* hybrids (tilapia) suggested a decrease of metabolism following direct exposure to glyphosate (Salbego *et al*., 2010; Muhammad *et al*., 2021)

Among metallic trace elements, the mercury (Hg), particularly under its bioaccumulative and toxic form methyl-mercury (MeHg), represents a major concern for ecosystems and human populations due to the high concentrations it reaches in predators (Lavoie *et al*., 2013; Kumsopar *et al*., 2025) and its neurotoxic effects (Chang, 1977). Fish exposed to Hg and/or MeHg present various biological effects (Zheng *et al*., 2019). For instance, Hg exposure can decrease hatching success of larvae, increase the embryo mortality of red seabream *Pagrus major* (Huang *et al*., 2011); and lead to skeletal deformities on larvae of *Danio rerio* (Liu *et al*., 2016). It can alter the oxidative stress response as observed on the gilt-head bream *Sparus aurata* and the tigerfish *Hoplias malaboricus* (Mela *et al*., 2014; Guardiola *et al*., 2016) and the immune response on goldfish *Carassius auratus* (Kong *et al*., 2012). Mercury exposure can affect the reproduction function as it disturbs hypothalamic hormones on *Danio rerio* and decrease the vitellogenin production on the fathead minnow *Pimephales promelas* (Klaper *et al*., 2006; Zhang *et al*., 2016). It can affect various organs, particularly the liver, causing damage and an increase in the hepatosomatic index in *Sparus aurata* (Guardiola *et al*., 2016).

Due to their toxicity, MeHg and glyphosate are considered as priority contaminants in terms of public health. Both contaminants were found almost everywhere in coral reef fish from south Pacific, including New Caledonia (Wejieme *et al*., 2025a; 2025b). This raises questions not only about their impact on the lagoon’s organisms but also about the human consumers who importantly rely on seafood for their subsistence. Among fish targeted for human consumption in SW Pacific and particularly in New Caledonia, the goldlined rabbitfish, *Siganus lineatus* (Valenciennes, 1835), is a tropical coastal reef fish found from mangroves to coral reefs with a high commercial value (Moleana, 2016). This fish is generally described as herbivore although Hoey *et al*. (2013) described this species as mixed feeders, feeding on a wide variety of algae, cyanobacteria, detritus and sediments. Herbivore fish are key component of reef ecosystems as they inhibit algal production and favoured coral implantation (Hughes *et al*., 2007). Due to their commercial value and to its role in ecosystems functioning, it is important to understand how fish are affected by these contaminants.

In this study, we investigated the influence of trophic contamination by MeHg and glyphosate on biology, physiology and sensorial system of *Siganus lineatus.* According to the literature, we predicted that these contaminants will influence negatively the body condition, growth and energy storage of fish. Through the study of the otolith’ shape (both *lapillus* and *sagitta*), we tested if the contaminants affect the vestibular system and therefore balance and/or hearing. We investigated if both *sagittae* in saccule and *lapilli* in utricle are affected in the same way.

## Materials and methods

### Diet contamination

Contaminants were incorporated onto floating pelleted feed (Le Gouessant) by coating using a DMSO/acidified water (30/70) solution. Four diets were prepared containing respectively 70 (MeHg 70), 700 (MeHg 700), 2100 (MeHg 2100) and 7000 (MeHg 7000) µg of organic mercury (Hg)/kg of feed with MeHg (TechLab); four other diets containing 100 (Gly 100), 1000 (Gly 1000), 3000 (Gly 3000), 10000 (Gly 10000) µg of glyphosate (TechLab) /kg of feed; and one diet (Control) was coated with the DMSO/acidified water solution without any contaminants.

### Experimental design

Three hundred rabbitfish, divided in two groups of 150 individuals, were transferred in 30 tanks (200 L-capacity), with 10 fish per tank. The first group used to assess the effect of MeHg measured 10.6 ± 1.1 cm in total length and weighted 32.2 ± 9.4 g wet weight; and 3 weeks later, the second group used to assess the effect of glyphosate measured 11.0 ± 1.2 cm and weighted 39.3 ± 11.9 g.

Fish of each group were acclimated to experimental conditions for four weeks before contamination experiment: temperature, water renewal, feeding sequence, daily interventions for water parameters monitoring, feeding and remains removal. Fish were fed with floating pelleted feed (Le Gouessant) during this acclimation period. Rations of 2-3% of biomass were distributed in two equal meals at 8 am and 2 pm. During the 2 hours meals, water renewal was stopped. Fish ate most of the feed immediately after distribution and the remains were removed after 2 hours, before restarting the water renewal.

The contamination phase lasted 7 weeks. In each group, fish were fed with 5 diets (3 tanks per diet): Control, MeHg 70, MeHg 700, MeHg 2100 and MeHg 7000 for the first experimental group; and Control, Gly 100, Gly 1000, Gly 3000, Gly 10000 for second one. Diets at 2.2 to 3 % of fish biomass were administrated in two equal meals as described for acclimation period. This feeding protocol was designed to limit contaminant (Hg or glyphosate) release from contaminated feed and ensure that exposure occurred mainly through the diet.

Physicochemical parameters of water were recorded daily, before feeding. The temperature was 27.8 ± 0.7°C, salinity of 33.6 ± 0.6, dissolved oxygen of 5.8 ± 0.5 mg L^−1^, and pH of 7.90 ± 0.08. Dissolved Hg levels in seawater source, determined before experimentation, were below detection limit. Dissolved contaminant possibly released from contaminated diet and by fish excretion was monitored with passive samplers for total Hg (DGT™ LSNB AP) and glyphosate (AFFINIMIP® POCIS), and was negligeable sources of Hg for fish.

### Fish handling and sampling

For all handling operations (fishing, transfer, weighing, measuring, sampling), the fish are handled under anaesthesia using eugenol diluted in 70° ethanol (0.2 mL of eugenol in 10 L of water). Once anesthetized, the fish can be handled quickly out of the water. During the transfer to the experimental room, the fish were individually marked by implantation of an electronic tag located in the dorsal muscle at the back of the head that can be read by a reader Biolog-id RS100 V8 (reference 9765). The fish were weighed and measured during the transfer to the experimental room (T_0_), then at the beginning (T_1_) and end of the experiment (T_2_), for individual monitoring of their growth. At the end of the experiment (T_2_), all fish were euthanized in a bath of eugenol comprising a dose 10 times greater than the anaesthesia dose (2 mL of eugenol in 10 L water), and kept on ice before dissection. Five fish per tank were dissected to extract and weight the liver.

### Influence on fish biology, physiology indices and contaminant levels

To evaluate the influence of contaminants on fish physiology, several biological indices such as Fulton’s condition factor, relative length and weight increases, weight growth rate, and hepatosomatic index were calculated (Table 1).

**Table 1.** Fish biological indices.

| Name | Formula | Explanation |
|---|---|---|
| Fulton’s condition factor (K) | $K = \frac{W}{TL^3}$ | where TL is the fish total length and W the fish weight |
| Weight growth rate (WGR) | $WGR = \frac{W_2 - W_1}{t_2 - t_1}$ | where W<sub>1</sub> and W<sub>2</sub> are the fish weight at time 1 et 2 |
| Relative length increase (RLI) | $\frac{TL_2 - TL_1}{TL_1}$ | where TL<sub>1</sub> and TL<sub>2</sub> are the fish total length at two different time |
| Relative weight increase (RWI) | $\frac{W_2 - W_1}{W_1}$ | where W<sub>1</sub> and W<sub>2</sub> are the fish weight at two different time |
| Specific growth rate (SGR) | $SGR = \frac{\ln W_2 - \ln W_0}{t_2 - t_0} \times 100$ | where W<sub>0</sub> and W<sub>1</sub> are the fish weight at time 0 et 2 |
| Hepatosomatic index (%) | $HSI = \frac{LW}{W} * 100$ | where LW is the liver weight and W the fish weight |

The Fulton’s condition factor explains the relation between length and weight of fish and give information on the global condition that can influence growth and survival. The relative length and weight increases informed on fish growth parameters.

The hepatosomatic index (HSI), the percentage of liver weight and fish mass ratio, was used as proxy of energy storage and to evaluate the effects of contaminants on fish (Muhammad *et al*., 2021). A decrease of HSI can be linked to a reduction in glycogen reserves due to increased energy expenditure in response to toxic exposure (Bonifacio *et al*., 2016) whereas an increase can be linked to pathological changes (hypertrophy or hyperplasia of hepatic cells).

### Influence on the sensorial system

As the morphology of both *sagitta* and *lapillus* can be implicated in sensorial systems (balance and hearing), the influence of contaminant on their shapes were studied. The otolith shape descriptors (Tuset *et al*., 2003) were used to describes the general aspect of otolith; and the elliptic Fourier descriptors for the tiny shape variations (e.g., Mérigot *et al*., 2007; Morat *et al*., 2012). After extraction and cleaning, both left and right *lapilli* and *sagittae* were digitised under a stereomicroscope (Leica I9) with the proximal face oriented toward the observer for *sagittae* and the ventral face for *lapilli* (Figure S1).

According to Tuset *et al*. (2003), three shape descriptors were determined from four otolith size parameters (area *A*, perimeter *P*, length *L* and width *W*). The form factor (*4*π*A/P²*) was used to estimate the irregularity of the surface area. This index ranges from 0 to 1, where 1 corresponds to a perfect circle. The roundness (*4A/*π*L²*) provides information on the similarity of various features to a perfect circle (index value of 1). The rectangularity (*A/LW*) informed on the variations in length and width of the otolith with respect to its area, which also ranges from 0 to 1 with 1 corresponding to a perfect square.

The tiny shape variation of the otolith contour was analysed by elliptic Fourier descriptors. The contour of otolith is described by components named harmonics, each harmonic is characterized by 4 coefficients resulting of the projection of each point of the contour on *x*- and *y*-axes. The higher the number of harmonics is, the greater the contour description is (Kuhl and Giardina, 1982; Morat *et al*., 2012). Each image of otolith was analysed with Shape v1.3 software that provide Fourier coefficients after make them invariants to the otolith size and its orientation (and position) regarding the beginning of the outline, which is arbitrarily defined (Iwata and Ukai, 2002). Regardless of the contaminant used, the Fourier power spectrum, determined through elliptic Fourier analysis (Crampton, 1995), showed that 99.98% of the outline reconstruction was obtained with 14 harmonics for *lapillus* and 14 for *sagitta* in average. The first harmonic is not taken into account (i.e., simple ellipse reconstruction), thus 52 Fourier coefficients were used to describe each otolith. Mean otolith shapes were reconstructed using *Momoc* R package using the same number of harmonics (Bonhomme *et al*., 2014).

### Statistical analysis

Anova and/or Kruskal-Wallis comparisons were used to evaluate the influence of contaminant concentrations on fish biological and otolith shape indices, depending on whether the data did fulfil the assumptions of normality and homoscedasticity.

Differences in otoliths shape (*sagittae* and *lapilli*) of *Siganus lineatus* exposed to various doses of contaminants were determined by canonical discriminant analysis (CDA) performed with Fourier coefficients. This classification method investigates the integrity of predefined groups (i.e., the different contaminant doses evoked in experimental design section) by finding a linear combination of the descriptors that maximise the Wilk’s lambda (λ) (Ramsay and Silveman, 2005). This statistic assesses the performance of the CDA, and it ranges from 0 (low discrimination) to 1 (high discrimination). Euclidian distances between barycentre of each group were calculated to quantify the dissimilarity between groups. A Cohen Kappa statistic was used to estimate the average proportion of individuals with index agreement, after removing the effect of chance agreement (Titus *et al*., 1984).

## Results

### Effect of contaminants on fish physiology

#### Mercury exposure

Survival rates were over 97% and fish grew from 10.59±1.12 cm for 32.25±9.39 g at T_0_ to 16.37±1.43 cm for 86.83±23.82 g at T_2_. The Fulton’s condition factor varied from 0.026±0.002 at T_0_ to 0.019±0.001 at T_2_. The relative length and weight increases (T_0_ to T_2_) were 0.55±0.13 and 1.80±0.73, respectively; whereas the specific growth rate was 1.29±0.33 %g *d*^−1^.

All biological and physiological indices (length and weight, weight growth rate, relative length and weight increased, specific growth rate, Fulton’s condition factor, and hepatosomatic index) did not show significant difference with methyl-mercury contamination (Tables 2 and 3).

**Table 2.** Mean (±sd) of *Siganus lineatus* parameters (length and weight) and fish biological indices measured before stabling (T_0_), at the beginning (T_1_) and at the end (T_2_) of contamination by methyl-mercury and glyphosate. W_0_, W_1_, W_2_, TL_0_, TL_1_, TL_2_ are the weight (g) and the total length (cm) of fish at T_0_, T_1_ and T_2_, respectively. K_0_, K_1_ and K_2_ are the Fulton’s condition factor (g cm^−3^) of fish at T_0_, T_1_ and T_2_. WGR_1_ and WGR_2_ are the weight growth rate (g *d*^−1^) at T_1_ and T_2_ respectively, RWI and RLI are the relative weight and length increase between T_0_ and T_2_. SGR is the specific growth rate (g *d*^−1^ in %) and the HSI is the hepatosomatic index (%) at T_2_. na is used when data are not available.

|  | Methyl-mercury |  |  |  |  | Glyphosate |  |  |  |  |
|---|---|---|---|---|---|---|---|---|---|---|
|  | Control | MeHg 70 | MeHg 700 | MeHg 2100 | MeHg 7000 | Control | Gly 100 | Gly 1000 | Gly 3000 | Gly 10000 |
| W<sub>0</sub> | 32.23±10.88 | 33.51±9.91 | 32.09±8.73 | 30.08±7.27 | 33.35±10.59 | 34.21±11.12 | 38.26±8.73 | 42.55±11.28 | 41.48±11.56 | 40.26±15.55 |
| W<sub>1</sub> | 45.08±13.87 | 49.10±13.45 | 47.56±11.75 | 45.21±10.25 | 49.92±15.38 | 46.05±14.29 | 52.58±13.00 | 56.26±11.73 | 54.52±14.90 | 53.45±17.04 |
| W<sub>2</sub> | 81.40±24.37 | 87.95±23.92 | 85.57±27.85 | 88.83±22.26 | 90.38±22.57 | 85.94±25.44 | 104.00±34.06 | 97.06±20.32 | 86.36±28.79 | 89.52±33.88 |
| TL<sub>0</sub> | 10.60±1.20 | 10.74±1.10 | 10.57±1.11 | 10.35±0.86 | 10.70±1.39 | 10.61±1.20 | 11.00±0.88 | 11.33±1.01 | 11.18±1.15 | 11.12±1.58 |
| TL<sub>1</sub> | na | na | na | na | na | na | na | na | na | na |
| TL<sub>2</sub> | 16.09±1.51 | 16.47±1.61 | 16.30±1.61 | 16.41±1.16 | 16.59±1.35 | 16.10±1.43 | 16.95±2.00 | 16.82±1.18 | 16.34±1.77 | 16.20±2.00 |
| K<sub>0</sub> (*10<sup>-2</sup>) | 2.63±0.17 | 2.64±0.15 | 2.65±0.10 | 2.67±0.18 | 2.64±0.19 | 2.78±0.20 | 2.84±0.16 | 2.87±0.19 | 2.91±0.22 | 2.83±0.19 |
| K<sub>1</sub> | na | na | na | na | na | na | na | na | na | na |
| K<sub>2</sub> (*10<sup>-2</sup>) | 1.90±0.13 | 1.92±0.09 | 1.91±0.14 | 1.98±0.12 | 1.94±0.09 | 2.00±0.18 | 2.04±0.09 | 2.02±0.12 | 1.91±0.20 | 2.01±0.10 |
| WGR<sub>1</sub> | 0.61±0.22 | 0.74±0.22 | 0.74±0.27 | 0.72±0.19 | 0.79±0.27 | 0.56±0.22 | 0.68±0.30 | 0.65±0.23 | 0.62±0.23 | 0.63±0.22 |
| WGR<sub>2</sub> | 0.65±0.28 | 0.69±.25 | 0.68±0.39 | 0.78±0.28 | 0.72±0.23 | 0.71±0.33 | 0.92±0.42 | 0.73±0.28 | 0.57±0.33 | 0.64±0.47 |
| RWI | 1.65±0.90 | 1.68±0.55 | 1.77±0.87 | 2.01±0.55 | 1.87±0.72 | 1.69±0.96 | 1.70±0.71 | 1.40±0.68 | 1.09±0.48 | 1.36±0.87 |
| RLI | 0.53±0.15 | 0.54±0.14 | 0.55±0.14 | 0.59±0.09 | 0.57±0.14 | 0.53±0.18 | 0.54±0.13 | 0.49±0.13 | 0.46±0.10 | 0.47±0.19 |
| SGR (%) | 1.21±0.37 | 1.25±0.30 | 1.26±0.41 | 1.41±0.26 | 1.33±0.31 | 1.21±0.46 | 1.26±0.37 | 1.09±0.36 | 0.92±0.31 | 1.03±0.51 |
| HSI (%) | 1.97±0.74 | 1.57±0.29 | 1.73±0.27 | 1.72±0.27 | 1.65±0.33 | 1.63±0.20 | 1.35±0.16 | 1.29±0.20 | 1.31±0.26 | 1.49±0.23 |

**Table 3.** Results of comparisons of contaminants effect on *Siganus lineatus* parameters and on fish biological indices at T_0_ (beginning of fish acclimation), T_1_ (beginning of contaminant exposure), T_2_ (end of contaminant exposure). F-statistic was used for Anova results and χ² for Kruskal-Wallis comparisons. Significative differences are shown in bold.

|  |  | Methyl-mercury | Glyphosate |
|---|---|---|---|
| Weight (W) |  |  |  |
|  | T<sub>0</sub> | F<sub>(4, 70)</sub> = 0.31, p = 0.87 | F<sub>(4, 70)</sub> = 1.15, p = 0.34 |
|  | T<sub>1</sub> | F<sub>(4, 70)</sub> = 0.43, p = 0.79 | F<sub>(4, 70)</sub> = 1.11, p = 0.36 |
|  | T<sub>2</sub> | F<sub>(4, 70)</sub> = 0.32, p = 0.87 | F<sub>(4, 70)</sub> = 1.08, p = 0.37 |
| Length (TL) |  |  |  |
|  | T<sub>0</sub> | χ<sup>2</sup> = 2.24, df = 4, p = 0.69 | F<sub>(4, 70)</sub> = 0.77, p = 0.55 |
|  | T<sub>1</sub> | na | na |
|  | T<sub>2</sub> | F<sub>(4, 70)</sub> = 0.25, p = 0.91 | F<sub>(4, 70)</sub> = 0.74, p = 0.56 |
| Fulton’s condition factor (K) |  |  |  |
|  | T<sub>0</sub> | F<sub>(4, 70)</sub> = 0.15, p = 0.96 | F<sub>(4, 70)</sub> = 0.97, p = 0.43 |
|  | T<sub>1</sub> | na | na |
|  | T<sub>2</sub> | χ<sup>2</sup> = 5.03, df = 4, p = 0.28 | F<sub>(4, 70)</sub> = 1.90, p = 0.12 |
| Weight growth rate (WGR) |  |  |  |
|  | At T<sub>1</sub> | F<sub>(4, 70)</sub> = 1.16, p = 0.34 | F<sub>(4, 70)</sub> = 0.48, p = 0.75 |
|  | At T<sub>2</sub> | F<sub>(4, 70)</sub> = 0.44, p = 0.78 | F<sub>(4, 70)</sub> = 1.84, p = 0.13 |
| Relative weight increase (RWI) |  |  |  |
|  | T<sub>0</sub> to T<sub>2</sub> | χ<sup>2</sup> = 4.99, df = 4, p = 0.29 | F<sub>(4, 70)</sub> = 1.71, p = 0.16 |
| Relative length increase (RLI) |  |  |  |
|  | T<sub>0</sub> to T<sub>2</sub> | F<sub>(4, 70)</sub> = 0.47, p = 0.75 | F<sub>(4, 70)</sub> = 0.78, p = 0.54 |
| Specific growth rate (%) (SGR) |  | F<sub>(4, 70)</sub> = 0.80, p = 0.52 | F<sub>(4, 70)</sub> = 1.55, p = 0.20 |
| Hepatosomatic index (%) (HSI) |  | χ<sup>2</sup> = 5.23, df = 4, p = 0.26 | **F<sub>(4, 70)</sub> = 6.77, p < 0.001** |

#### Glyphosate exposure

Fish survival rates ranged from 83% (Gly 3000) to 100% (Gly 100 and Gly 1000). They grew from 11.05±1.18 cm for 39.35±11.90 g at T_0_ to 16.48±1.70 cm for 92.58±29.04 g at T_2_. The Fulton’s condition factor varied from 0.028±0.002 at T_0_ to 0.020±0.001 at T_2_. The relative length and weight increases (T_0_ to T_2_) were 0.50±0.15 and 1.45±0.77, respectively; although the specific growth rate was 1.10±0.41 % g *d*^−1^.

All biological and physiological indices (length and weight, weight growth rate, relative length and weight increased, specific growth rate, and Fulton’s condition factor) did not show significant difference with glyphosate contamination (Tables 2 and 3), except the hepatosomatic index which was significantly lower for fish contaminated by glyphosate (Figure 1).

**Figure 1.**
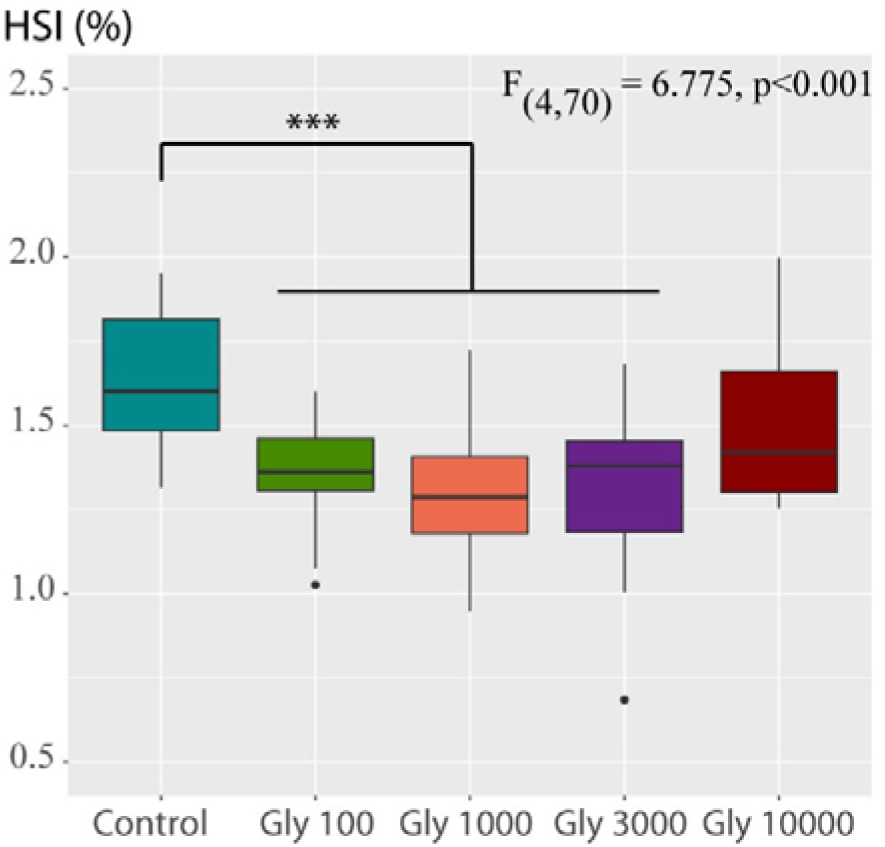
Boxplot of hepatosomatic index in function of glyphosate (Gly) concentrations in diet. F-statistic was used for Anova results. * p < 0.05, ** p < 0.01.

### Effect of contaminants on sensorial system

#### Methyl-mercury exposure

The general aspect of otolith measured by the shape indices differed only for *lapilli* and not for *sagittae* (Figures S2 and S3). The form factor and the roundness were lower for fish contaminated with high concentration of MeHg (MeHg 7000) than control for left *lapilli* whereas these indices were lower for fish contaminated with low concentration of MeHg (MeHg 70) than control for right *lapilli*. Rectangularity of both *lapilli* was not affected by MeHg contamination.

The tiny description of the otoliths outline based on Fourier coefficients showed a clear discrimination for all treatments for both *lapilli* (Figure 2A, B) and *sagittae* (Figure 2C, D) with high reclassification by Cohen-Kappa (96.5 – 100%). While discriminations based on *lapilli* shapes did not show dose effect organisation (Figure 2A, B), the *sagittae* showed a discrimination with dose effect on the first discriminant function for both otolith side (Figure 2C, D). *Sagittae* from fish from Control and those contaminated by low doses of MeHg (MeHg 70 and MeHg 700) were clearly separated from those exposed to high doses (MeHg 2100 and MeHg 7000). For all discriminant analysis, all groups were separated with high Euclidian distance between centroids (Table S1).

**Figure 2.**
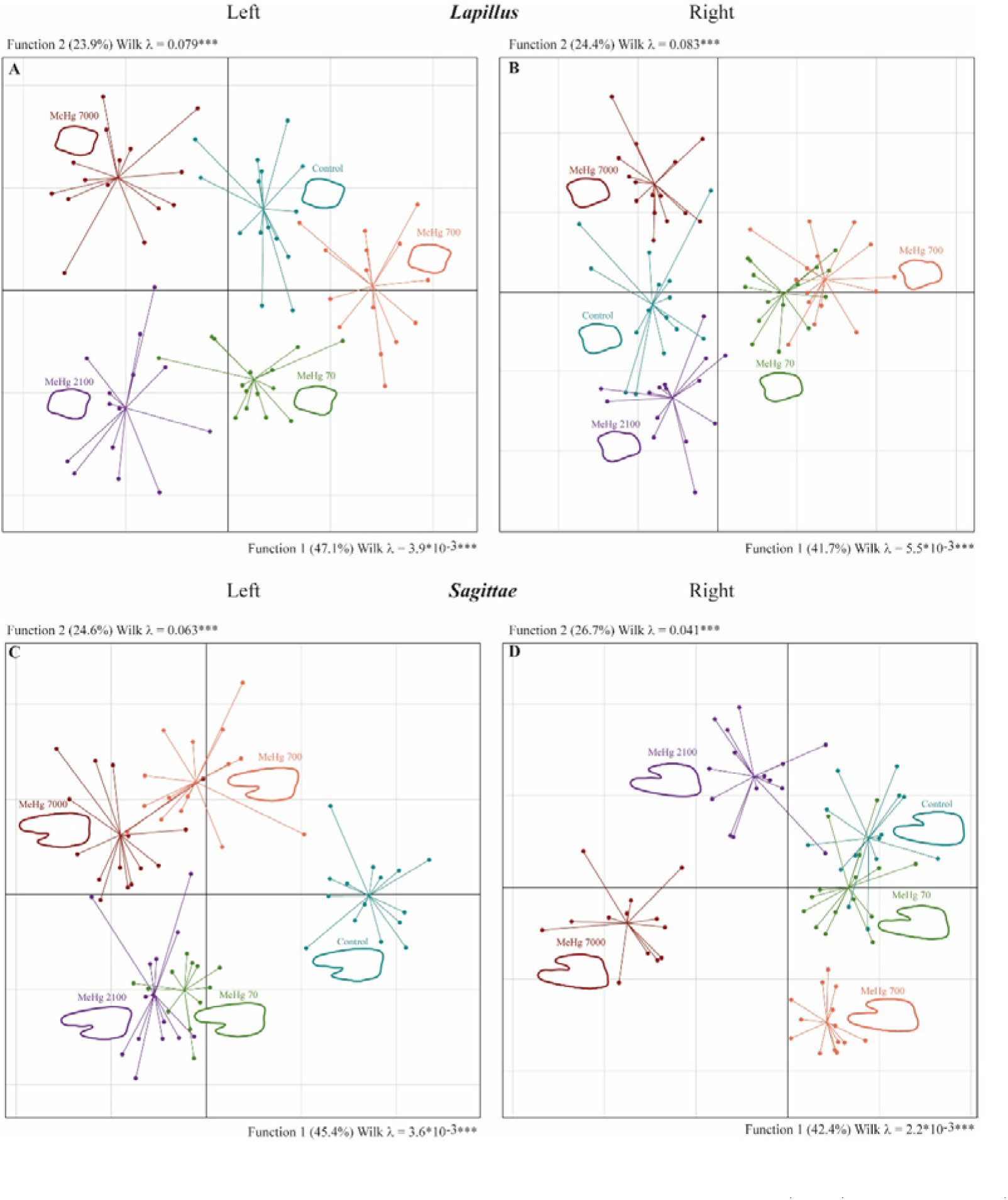
Discriminant analyses performed with Fourier coefficients on *lapilli* (A, B) and *sagittae* (C, D) of *Siganus lineatus* exposed to various doses of methyl-mercury (MeHg). Average otolith of groups was represents using *Momoc* R-parckage.

The *lapillus* average shapes of fish exposed to MeHg doses showed more variations for left than right side (Figure 3). Left *lapillus* varied all around the contour of otolith with stronger variations on anterior and posterior faces whereas variation on right side were tinier. Variations on *sagittae* were similar for both sides and were located on notch area and on anterior (*rostrum*) and posterior faces (*post-rostrum*) (see Figure S1 for otolith description).

**Figure 3.**
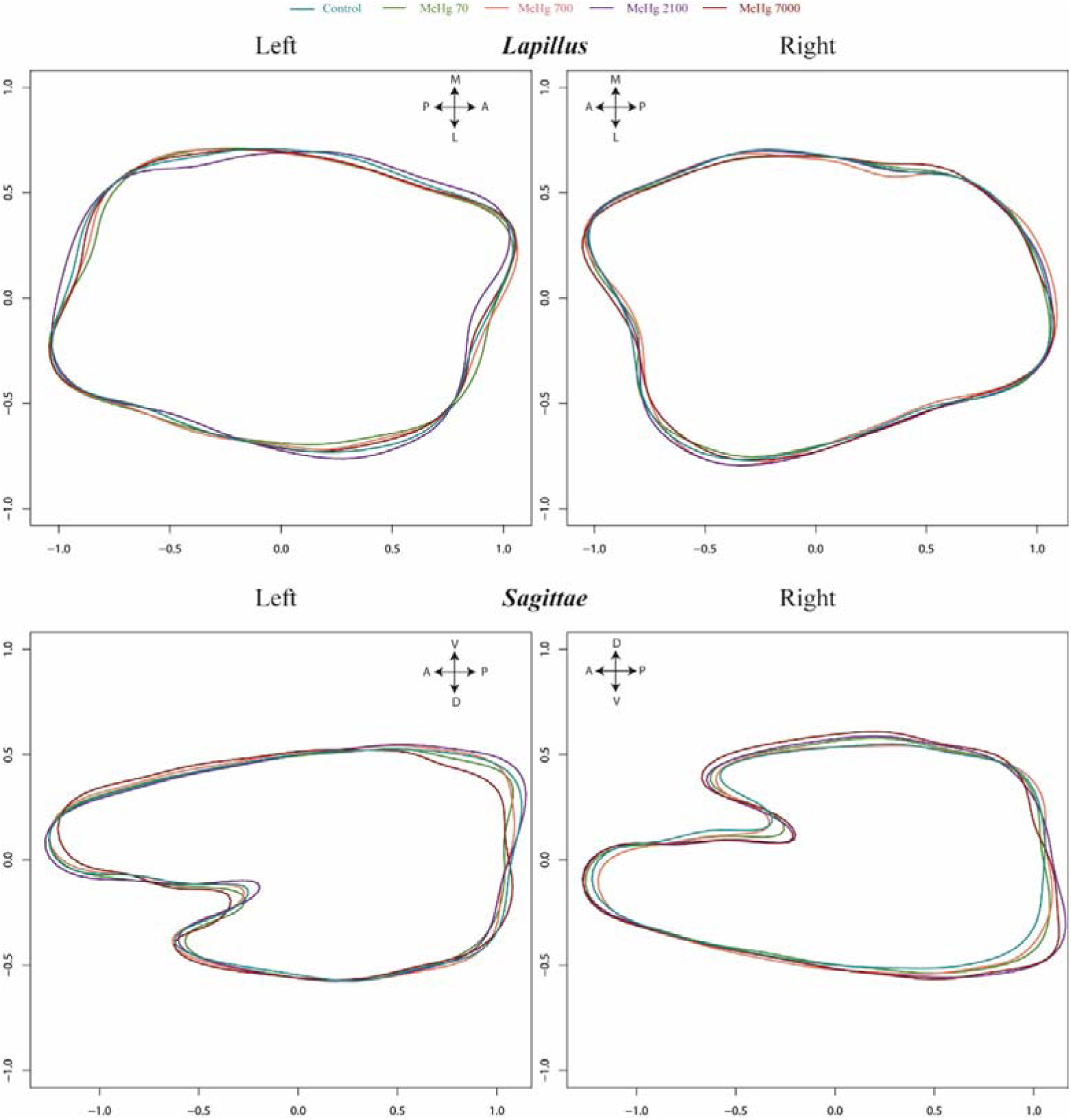
Reconstruction of the average *lapilli* and *sagittae* of *Siganus lineatus* exposed to various doses of methyl-mercury (MeHg). A = anterior face, D = dorsal face, M = Medial (proximal) face, L = lateral (distal) face, P = posterior face, V = ventral face.

### Glyphosate exposure

The general aspect of otolith measured by the shape indices differed only for *lapilli* and not for *sagittae* (Figures S2 and S3). Form factors were higher for fish contaminated with high concentration of glyphosate (Gly 1000, Gly 3000 and Gly 10000) than control and Gly 100 for both left and right *lapilli*. Rectangularity of both *lapilli* was not affected by glyphosate contamination.

The tiny description of the otoliths outline based on Fourier coefficients showed a clear discrimination for all treatments for both *lapilli* (Figure 4A, B) and *sagittae* (Figure 4C, D) with high reclassification by Cohen-Kappa (90.0 – 100%) with high Euclidian distances (Table S1). No clear dose effect organisation was observed except for right *lapilli* and left *sagittae* where Control and low glyphosate (Gly 100) contamination were separated from high doses (Figure 4B, C).

**Figure 4.**
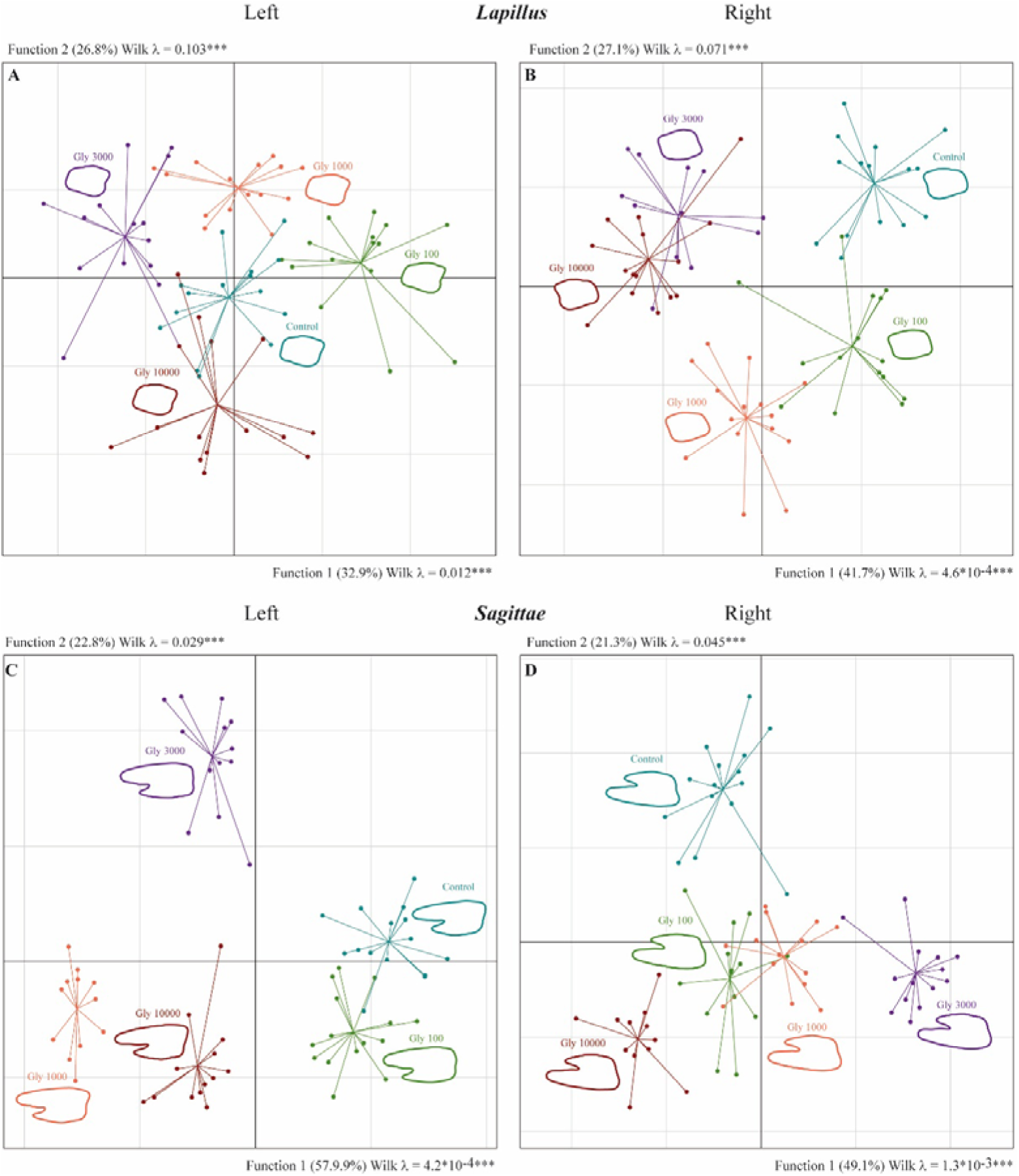
Discriminant analyses performed with Fourier coefficients *lapilli* (A, B) and *sagittae* (C, D) of *Siganus lineatus* exposed to various doses of glyphosate (Gly). Average otolith of groups was represents using *Momoc* R-parckage.

The *lapillus* average shapes of fish exposed to glyphosate doses showed variations for both left and right sides (Figure 5). *Lapillus* varied all around the contour of otolith with stronger variations on anterior (*gibbus maculae*) and posterior faces. Variations on *sagittae* were stronger for left than right side and were located on notch area, on anti-rostrum and posterior face (*post-rostrum*).

**Figure 5.**
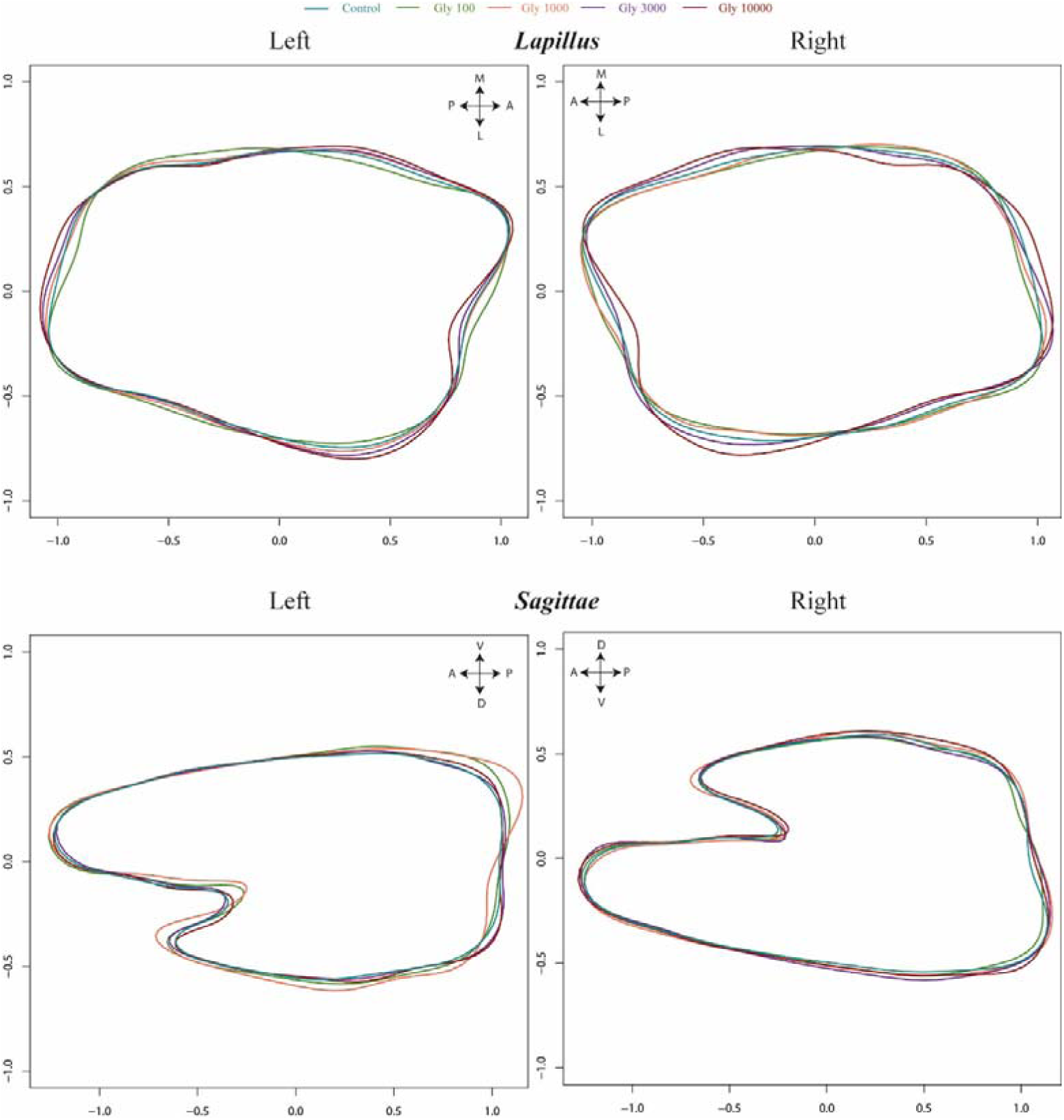
Reconstruction of the average *lapilli* and *sagittae* of *Siganus lineatus* exposed to various doses of glyphosate (Gly). A = anterior face, D = dorsal face, M = Medial (proximal) face, L = lateral (distal) face, P = posterior face, V = ventral face.

## Discussion

Our study shows that *Siganus lineatus* exposed to MeHg or glyphosate presented only few physiological alterations (lower hepatosomatic index for glyphosate and for the highest concentrations of both contaminants in muscle. However, both contaminants had modified the shape of both *lapillus* in utricle and *sagitta* in the saccule and so can have impacted the sensorial system.

### Biological responses to contaminants

Contrary to our hypothesis, the biological parameters were not negatively affected by contaminants (except for glyphosate on HSI). Influence of contaminants on HSI and fish condition are contrasted in the literature. For example, European seabass *Dicentrarchus labrax* and gilthead seabream *Sparus aurata* exposed to MeHg showed an increase of HSI (Guardiola *et al*., 2016; Maulvault *et al*., 2017) due to an increase of the enzymatic activity responsible of the detoxification of contaminants, while the ten spotted live-bearer (*Cnesterodon decemmaculatus*) and crossbred red tilapia (hybrid of *Oreochromis niloticus* and *O. mossambicus*) exposed to glyphosate showed no effect on HSI (Bonifacio *et al*., 2016; Muhammad *et al*., 2021). The fish condition decreased for European seabass and ten spotted live-bearer exposed to MeHg and chlorpyrifos (another pesticide) respectively, whereas it was not affected for crossbred red tilapia and *Leporinus obtusidens* exposed to glyphosate (Salbego *et al*., 2010; Bonifacio *et al*., 2016; Maulvault *et al*., 2017; Muhammad *et al*., 2021).

In this study, fish exposed to contaminant did not showed difference in growth or body condition. In both experiments, the fish were fed at 2-3% of the biomass; this food and therefore energy supply could allow them to respond more effectively to contamination by supporting the energetic cost of the detoxification processes. Similarly, Carravieri *et al*. (2022) on a bird meta-analyse showed no impact of Hg on body condition, particularly when birds are fed *ad libitum* in controlled conditions. The hepatosomatic index is a more specific biomarker than Fulton’s condition factor as it is directly linked to toxic effects of contaminant on liver (Bonifacio *et al*., 2016). The small decrease of HSI with glyphosate exposure could indicate a depletion of glycogen reserves due to an increase of energetic expenditure in response to the toxic event. The absence of effect observed with the MeHg exposure could reflect either a depletion of glycogen or a pathological change such as hyperplasia of hepatic cells and/or liver hypertrophy (Bonifacio *et al*., 2016).

### Effect of contaminants on sensorial system

Both studied otoliths of *S. lineatus* exposed to contaminants showed shape variations, supporting our hypothesis of potential effects of contaminants on the vestibular system and therefore balance and/or hearing. *Sagittae* and *lapilli* varied mainly in antero-posterior axis, with stronger variations on *rostrum*, *anti-rostrum*, *post-rostrum* and notch area for *sagittae* and in *gibbus maculae* area for *lapilli*. Several studies have evidenced the shape variations of otolith with environmental characteristics (temperature, habitats, disturbances, etc.), however only few had studied the influence of contaminants on otolith shape. In Finland, burbots (*Lota lota*) from a polluted lake by metals and PCBs presented otoliths with crenulations absent in other populations (Pulliainen and Korhonen, 1994). Based on *sagitta* analyses, an annular sea bream (*Diplodus annularis*) population from contaminated area (Cr, Fe, Pb, and Zn) was discriminated from those in uncontaminated area in Tunisia (Gharred *et al*., 2020). Bose *et al*. (2018) showed that round goby (*Neogobius melanostomus*) from contaminated areas had *sagittae* with more pronounced caudal points (equal to *post-rostrum*) and less prominent ventral protrusions. In a recent study, Beauvieux *et al*. (2026) showed that *sagittae* of gilthead seabream (*Sparus aurata*) evidence shape differences with pollutants in three coastal lagoons in Tunisia. They evidenced that shape variations were found predominantly in left *sagittae* and were associated with cupper and rubidium; as we have observed for MeHg contamination on our study. Contrary to our study in controlled laboratory conditions, field studies may be affected by confounding habitat characteristics that can act on otolith shape.

Investigations on the influence of shape on otolith function are scarce but fish with abnormal otoliths (malformations) showed a disturbed schooling behaviour and no response to sound stimuli (Browning *et al*., 2012), as well as altered escape movements (Vignon and Aymes, 2020). Schulz-Mirbach *et al*. (2019) suggested that otolith shape and mass could influence the inner ear function on two main ways. Firstly, otoliths could move on a simple manner (back- and-forth) along the rostro-caudal or dorso-ventral axes when the fish swims, tilts, or during left to right movements (detected by *lapillus* that is horizontally oriented, Fig. S1, Assis (2005)). The positional changes and/or the movement of fish would generate a linear acceleration that could induce the otolith motion detected by the sensory epithelium (*macula)*. In this case, the otolith motion would depend mainly on their mass, but the different otolith shapes could induce different flow dynamics of the endolymphatic fluid when the otolith moves relative to the sensory epithelium. Secondly, if the movement/oscillation of the otolith is induced by sound stimulus generating an oscillating motion field over the entire fish body, the oscillation pattern could be a simple back-and-forth movement relative to the sensory epithelium if the otolith is roughly spherical or ellipsoidal but could deviate for more complex shape (with humps or flutes). Gauldie and Crampton (2002) evidenced that fish from deep environments had more complex otolith shape’s that could increase their hearing; an important sense in dark environment for detecting prey and avoid predators.

In our study, the otolith shape of *S. lineatus* varied in the notch area for *sagitta* (opening of *sulcus acusticus*) and in *gibbus maculae* area for *lapilli*. The s*ulcus* and the *gibbus* have an important role as they are directly in contact with the sensory epithelium (*macula*) (Assis, 2005; Tuset *et al*., 2008), and their shapes closely matches their respective *macula*. It is assumed that the ratio of otolith area to the *sulcus* area results to a modification of hearing abilities (alteration or improve) (Ramcharitar *et al*., 2004; Schulz-Mirbach *et al*., 2010; 2019). The potential effect of the size or shape of the *sulcus* or *macula* to those of otoliths are not experimentally tested, however we can suggest that they can impact on the hearing ability or equilibrium. For fish contaminated by MeHg and glyphosate, the variability in the opening area of *sulcus* and *gibbus* could suggested that contaminants affect the functions of inner ear.

### Are both otoliths affected in the same way?

In the review on the functional role of otolith, Schulz-Mirbach *et al*. (2019) question about if the three otoliths are sensitive in the same manner to environmental variations. In the present study, the shape of both otoliths formed on the beginning of ontogenesis (*sagittae* and *lapilli*) (Lecomte-Finiger, 1999; Bounket *et al*., 2019) was studied, whereas all other studies used only one single otolith (mainly the *sagitta*). Both otoliths of *S. lineatus* are affected by contaminants; in the MeHg experience the *sagittae* is affected whereas it was the *lapillus* in the glyphosate experience. This could suggest that both otoliths contained in saccule and utricle are not affected in the same way. Despites that we are in experimental conditions with higher contaminant concentrations in most experimental cases than in natural environment, we showed that a single contaminant is able to influence otoliths shape. In the environment, contaminants are never alone but are part of a group of compounds that can sometimes be numerous, and this mixture (‘cocktail effect’) could have addictive and/or synergetic effects on vestibular system impacting both equilibrium, motion perception and audition.

## Acknowledgments

The authors thank the French “Agence nationale de la recherche” for founding the TONIC project (ANR-21-CE32-0008).

## Contributions

YL, FM, NW, ET, VP, and PB conceived the project. NW, CL, FG, MM and FA performed the experimental experience from the contamination of the diet to the fish rearing. NW, YL and FM dissected fishes. FM performed the analyses and prepared the figures. FM, NW, ET, VP, PB and YL drafted the main text of the manuscript. All authors revised and approved the manuscript.

## Data availability

Raw data used in this study are available in SEANOE repository by the link https://doi.org/10.17882/121532 (Morat *et al*., 2026).

