## Supplementary material for "Effects of an experimental contamination by methyl-mercury and glyphosate on biological and otolith shape parameters of a tropical Siganidae": Suplemental tables and figures

Table S1. Euclidian distances between groups centroid of canonical discriminant analyses achieved with *lapilli* and *sagittae* of *Siganus lineatus* contaminated with methyl-mercury (MeHg) and glyphosate (Gly).

| **Methyl-mercury** | | | | | | |
| --- | --- | --- | --- | --- | --- | --- |
| ***Lapillus*** |  |  | MeHg 70 | MeHg 700 | MeHg 2100 | MeHg 7000 |
|  | Control | Left  Right | 1.67  1.65 | 1.31  2.19 | 2.36  1.20 | 1.45  1.52 |
|  | MeHg 70 | Left  Right |  | 1.47  0.55 | 1.28  1.91 | 2.37  2.13 |
|  | MeHg 700 | Left  Right |  |  | 2.69  2.42 | 2.70  2.46 |
|  | MeHg 2100 | Left  Right |  |  |  | 2.25  2.70 |
| ***Sagittae*** | Control | Left  Right | 2.18  0.58 | 2.18  2.06 | 2.48  1.42 | 2.69  2.79 |
|  | MeHg 70 | Left  Right |  | 2.20  1.49 | 0.32  1.59 | 1.77  2.45 |
|  | MeHg 700 | Left  Right |  |  | 2.28  2.80 | 0.97  2.43 |
|  | MeHg 2100 | Left  Right |  |  |  | 1.72  2.12 |
| **Glyphosate** | | | | | | |
| ***Lapillus*** |  |  | Gly 100 | Gly 1000 | Gly 3000 | Gly 10000 |
|  | Control | Left  Right | 1.55  1.66 | 1.25  2.70 | 1.36  2.00 | 1.23  2.40 |
|  | Gly 100 | Left  Right |  | 1.64  1.29 | 2.69  2.20 | 2.29  2.24 |
|  | Gly 1000 | Left  Right |  |  | 1.40  2.16 | 2.48  1.88 |
|  | Gly 3000 | Left  Right |  |  |  | 2.18  0.53 |
| ***Sagittae*** | Control | Left  Right | 0.84  2.00 | 2.76  1.88 | 2.22  2.81 | 1.97  2..78 |
|  | Gly 100 | Left  Right |  | 2.40  0.61 | 2.68  1.96 | 1.38  1.16 |
|  | Gly 1000 | Left  Right |  |  | 2.48  1.41 | 1.15  1.77 |
|  | Gly 3000 | Left  Right |  |  |  | 2.68  3.01 |

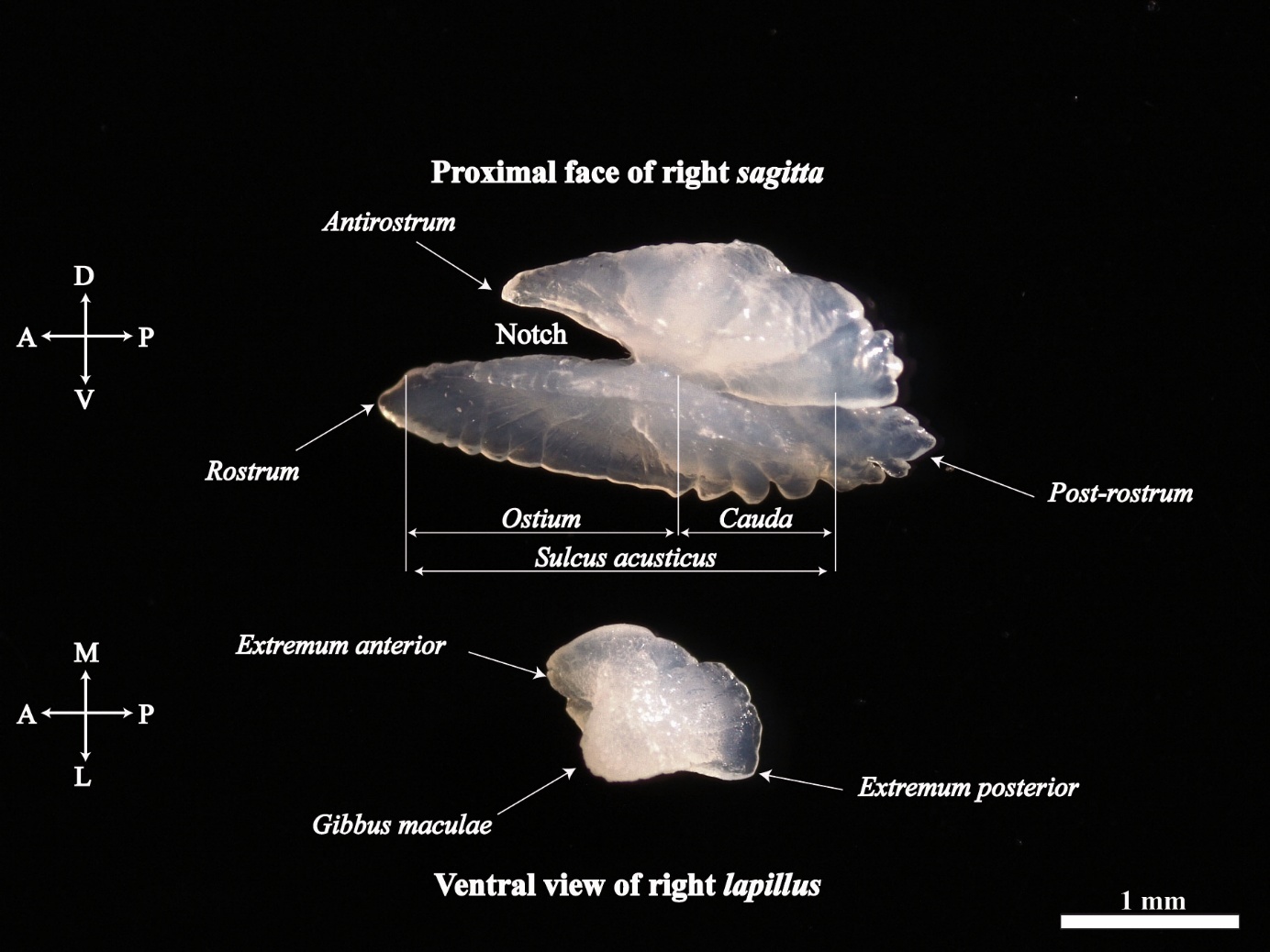

Figure S1. Annotated image of right *sagitta* and *lapillus* of *Siganus lineatus* digitised on proximal and ventral views, respectively. Terminology are in accordance with Tuset *et al.* (2008) for *sagitta* and Assis (2005) for *lapillus*. A = anterior face, P = posterior face, D = dorsal face, V = ventral face, M = Medial (proximal) face, L = lateral (distal) face.

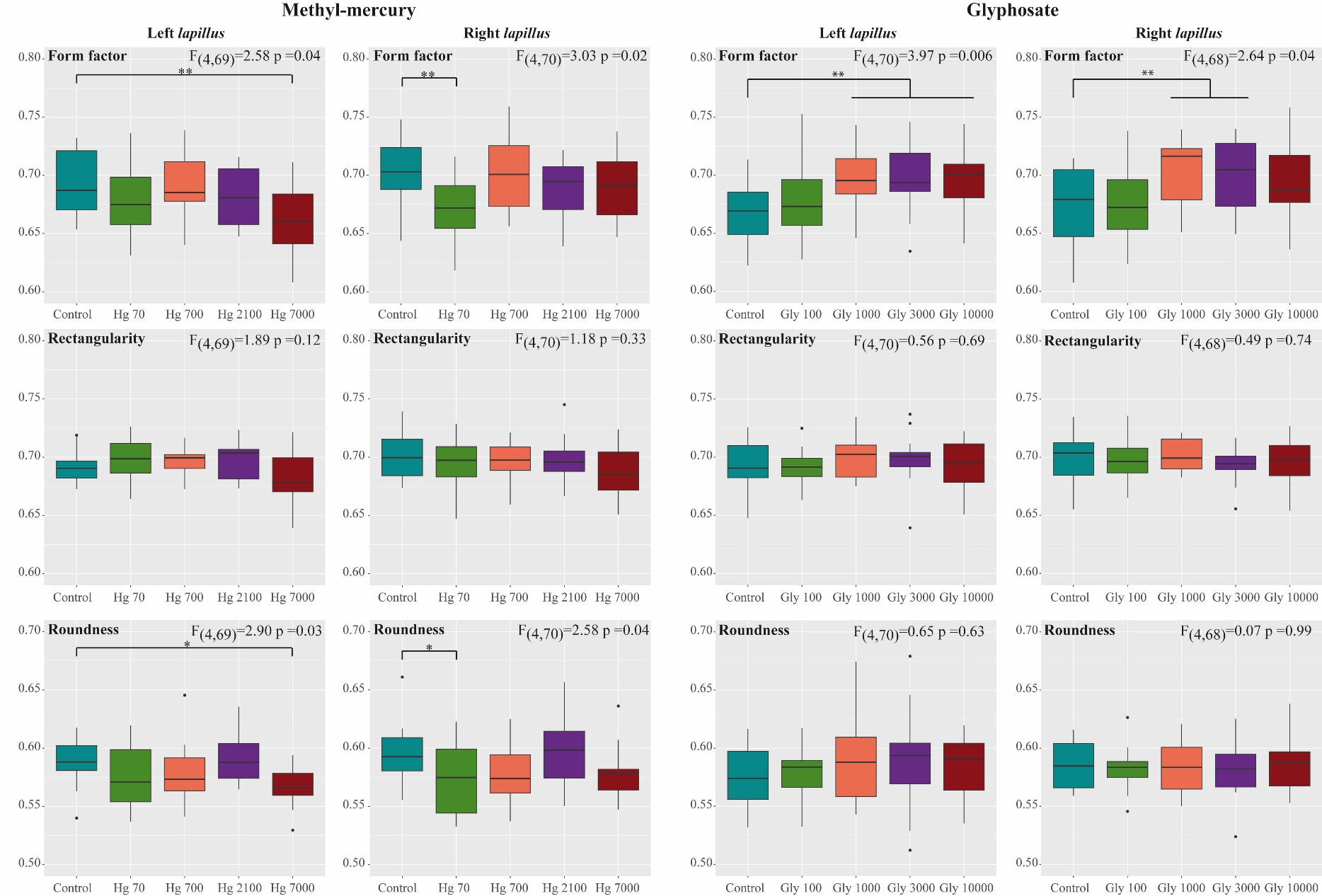

Figure S2. Boxplot of l*apilli* shape indices of *Siganus lineatus* in function of contaminants concentrations in diet. Hg = methyl-mercury and Gly = glyphosate. F-statistic was used for Anova results. * p < 0.05, ** p < 0.01.

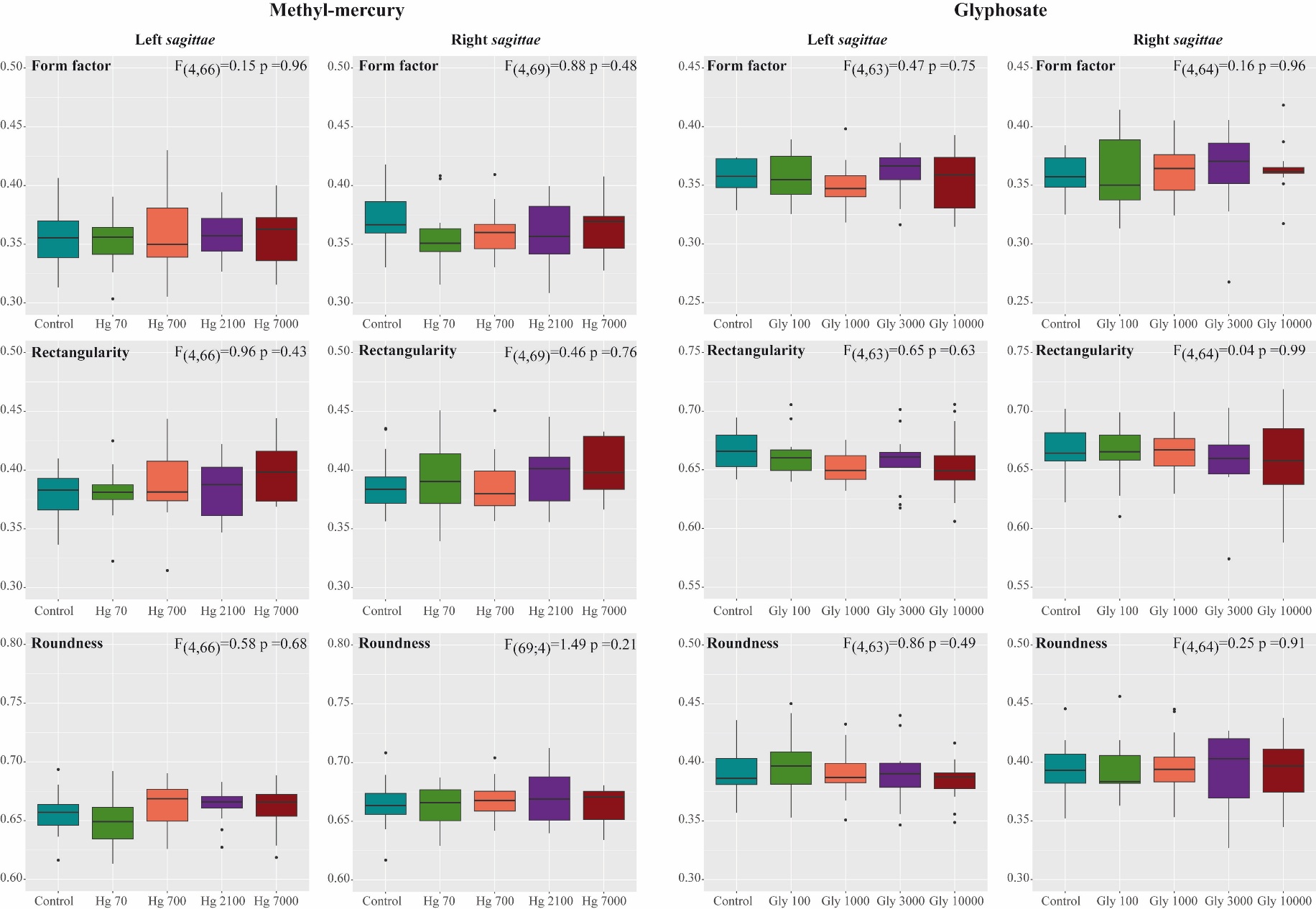

Figure S3. Boxplot of *sagittae* shape indices indices of *Siganus lineatus* in function of pollutants concentrations in diet. Hg = methyl-mercury and Gly = glyphosate. F-statistic was used for Anova results. * p < 0.05, ** p < 0.01.
